# Improving drug safety predictions by reducing poor analytical practices

**DOI:** 10.1101/2020.09.25.314138

**Authors:** Stanley E. Lazic, Dominic P. Williams

## Abstract

Predicting the safety of a drug from preclinical data is a major challenge in drug discovery, and progressing an unsafe compound into the clinic puts patients at risk and wastes resources. In drug safety pharmacology and related fields, methods and analytical decisions known to provide poor predictions are common and include creating arbitrary thresholds, binning continuous values, giving all assays equal weight, and multiple reuse of information. In addition, the metrics used to evaluate models often omit important criteria and models’ performance on new data are often not assessed rigorously. Prediction models with these problems are unlikely to perform well, and published models suffer from many of these issues. We describe these problems in detail, demonstrate their negative consequences, and propose simple solutions that are standard in other disciplines where predictive modelling is used.

## Introduction

In early-stage drug discovery, compounds are tested in assays to determine their likelihood of causing organ toxicity or clinical adverse events. The assays often test specific mechanisms such as blocking key ion channels, inhibiting mitochondrial function, causing cell death, or damaging DNA. Based on these assay results and other information about the compounds such as their structural, physical, and chemical properties, project teams decide whether to progress a compound to animal studies, and eventually to human trials. At each stage of drug discovery pipeline, decisions are made based on current data, and quantitative methods are developed to assist decision-makers [1].

Many current practices are statistically inefficient and misleading. The aim of this paper is to describe these practices, illustrate their limitations, and demonstrate the benefits of alternative approaches using simple examples and simulations. These recommendations complement existing guidelines and best practices on *in silico* toxicology models [2, 3].

## Problems with current practices

### Using safety margins as predictors

Compounds are often tested in concentration-response assays and the results are typically summarised by an XC_50_ value, which is a measure of the potency of the compound, and which lies between the lowest and highest concentrations tested. Another important quantity for safety assessment is the peak or maximum concentration that a compound reaches in the blood (C_max_) – either actual or predicted. C_max_ is related to the clinical dose, and compounds given at higher doses tend to have a greater risk for toxic side effects, making C_max_ one of the best predictors of clinical toxicity [4, 5, 6]. XC_50_ values are often divided by C_max_ values to give a safety margin or safety factor, where higher values indicate greater safety. Although safety margins are useful and easy to interpret, they have five problems when used to make predictions. The first problem is minor: an assumption is created that all combinations of XC_50_ and C_max_ values with the same margin have the same risk. For example, 1/0.1, 10/1, 100/10 all have a margin of 10, and therefore are assumed to have the same risk. This may be a reasonable assumption, but it should be verified.

A more damaging consequence of using margins is that mechanistic assays with zero predictive ability for clinical toxicity can appear useful. For example, mechanistic assays are used to design out specific liabilities such as mitochondria inhibition or bile salt export pump inhibition. Even if the primary purpose of these assays is for drug design, they are often used in models to predict clinical toxicity, but the margin values are used instead of the XC_50_ values. Using all available data is a good idea, but we show in Figure 1 with simulated data how margins can mislead. XC_50_ values for two assays are independently simulated from a uniform distribution for 100 compounds, and an outcome of clinical interest is also independently simulated from a uniform distribution. Figure 1A and 1B show no association between the assay values and the outcome, indicating that the assays have zero predictive ability. Next we simulate C_max_ values that strongly predict the outcome (Fig. 1C), and then calculate the safety margins. Both margins are now good predictors of the outcome (Fig. 1D and 1E), but this is driven entirely by C_max_, the assays have contributed nothing to the prediction.

**Figure 1.**
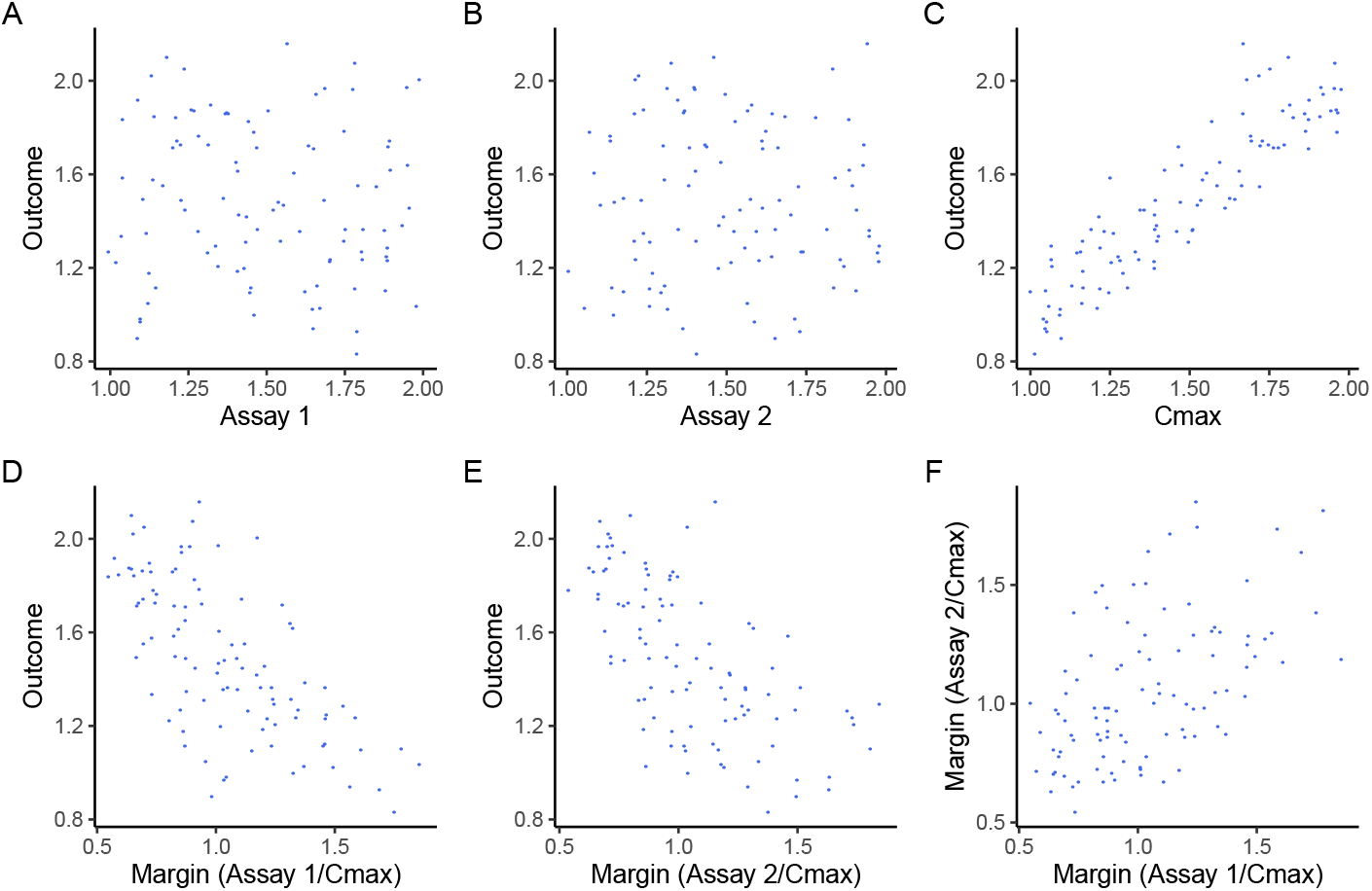
Using safety margins can mislead. Neither Assay 1 (A) nor Assay 2 (B) predict the outcome, but C_max_ is strongly predictive (C). Safety margins for Assay 1 (D) and Assay 2 (E) appear predictive, but these relationships are driven entirely by C_max_. The assay results are now correlated, making prediction models less stable (F).

A third problem is that the safety margin variables are now correlated, owing to their dependence on C_max_ (Fig. 1F), whereas the original assay values are uncorrelated (not shown, but the values were simulated to be independent). Correlated variables are a problem when creating prediction models because the models become less stable, the precision of the estimates decreases, and the apparent importance of the variables can decrease.

A fourth problem is that the predictive information contained in the C_max_ values is used multiple times and thus C_max_ receives greater weight. Even if the assays are predictive, C_max_ is one piece of information that should be used only once, not multiple times for each margin. Furthermore, C_max_ is often measured with considerable uncertainty due to variability between patients, or it may be predicted from *in vitro* and *in silico* data. This measurement noise is then added to all assay values.

Finally, by combining two variables into one, we’ve reduced our flexibility to make predictions. For example, Figure 1A in Lazic et al. plots margin values for a cardiotoxicity model, and the only option to separate the two classes of compounds is the location of a vertical line [5]. Figure 1B in that paper plots C_max_ and the IC_50_ values from a hERG inhibition assay on two axes, and now it is easier to separate the groups with a line in two dimensions, as both the slope and intercept provide greater flexibility. Furthermore, we have even more flexibility because the separating boundary need not be a straight line – a curved line might be preferable. Reducing the dimensions of the data by calculating a ratio of two values can be beneficial when faced with too many predictor variables, but the trade-off is reduced flexibility. Given all these problems with safety margins, they will rarely be good inputs into a predictive model. Instead, using C_max_ and XC_50_ values directly will often be preferable.

### Binning and arbitrary thresholds

A common practice is to categorise compounds into “Active” or “Not active” in an assay based on their XC_50_ values or safety margins. This makes as much scientific sense as measuring people’s height with a noisy ruler and then categorising them into “giants” and “dwarfs” based on an arbitrary cutoff. There is little gained by such a procedure [7, 8, 9, 10, 11, 12, 13, 14, 15, 16, 17].

XC_50_ values, like human height, are continuous variables (at least to the measured precision) and should be treated as such. Binning compounds into Active/Not active has three problems. First, information is discarded. All compounds within a category are considered equal, but scientific sense tells us that a compound just below a threshold will differ from another compound in the same category that is far from the threshold. Similarly, compounds just on either side of a threshold usually do not have dramatically different risks. Second, binning creates categories that do not exist in nature, but are then taken as real. This is known as the reification fallacy, or the fallacy of misplaced concreteness, where a concept is mistaken for a physical property. This leads to surprise when compounds that are “negative” in all assays show clinical toxicity later on. Being negative in an assay is not a physical property of a compound, it’s an arbitrary designation we’ve assigned. A compound with a weak signal in several assays may be deemed safe because the XC_50_ values were below a threshold, but this does not reflect the true risk. An additive or multiplicative combination of several weak signals could potentially indicate a high-risk compound.

Third, binning ignores uncertainty in the assay values; 95% confidence intervals for XC_50_ values can span several orders of magnitude. If an assay was re-run, compounds close to the threshold could fall on the opposite side (Elkins et al. discuss variability in measured IC_50_ values and how to account for them in computational models [18]). This procedure therefore introduces misclassification errors in the data. Thresholds are only needed when a decision or action is taken, which occurs after a prediction is made (or based on assay values directly if no prediction is made; for example, selecting compounds to progress from a high-throughput screen). When building predictive models, the numeric assay values should be used.

### Deriving arbitrary “scores”

To rank compounds or make a decision, the information contained in multiple variables that predict clinical safety need to be combined. These variables are often on different scales and are not directly comparable; for example, assay results are often XC_50_ values, physical and chemical properties such as cLogP may have no units, and compounds satisfying Lipinski’s Rule-of-5 may be binary Yes/No variables. One motivation for binning continuous variables is that all become binary and can be combined into a score by summing them, but ensuring that the value of “1” is interpreted as “higher risk” for all variables:

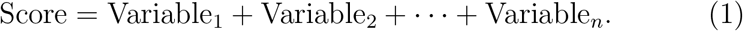

The score can range from zero to the number of variables n, and the hope is that the score predicts the clinical outcome. We refer to this as the “bin-and-sum” method, which was previously used at AstraZeneca and other companies [19, 20]. Fortunately, the above score equation is the beginning of a standard statistical model. In Equation 2 below we convert the above score equation into a statistical model by using the measured variable values directly to predict the clinical outcome *y*, which could be QT_c_ prolongation for a cardiotoxicity model or liver enzyme levels for a liver toxicity model (we assume y is a continuous variable for this discussion, but the equation below can be modified for other types of outcomes). We also introduce parameters (*β*’s, which are called weights in the machine learning literature) that are estimated from the data. The above bin-and-sum method also implicitly has parameters, but they are fixed to *β*_0_ = 0 and the other β’s to a value of 1, and therefore have no influence.

Why fix the *β*’s? It is better to learn their optimal values from the data. Note how in the equation below we are using the variables to directly predict the outcome y, not to sum to a score which we hope will be associated with the outcome in a later step. Also, the variables do not need to be on the same scale or have the same units.

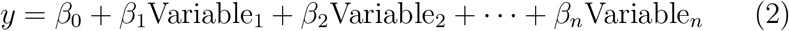

To complete the statistical model we also need an error term (*ε*), which captures the difference between the prediction based on the variables and the true value of the outcome:

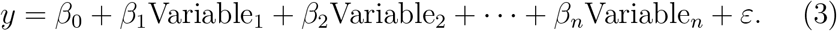

The bin-and-sum method has several problems. First, the predictor variables are binned, which was discussed in the previous section. Second, all variables are given equal weight in calculating the score, implying that all variables are equally effective in predicting the outcome, but this is unrealistic. Third, the outcome – the target of our prediction – is not used to create the prediction rule, which is an inefficient use of the data. Fourth, even simple interactions between variables cannot be captured. Figure 2 illustrates this point with simulated data. 100 compounds are defined as either toxic (red triangles) or safe (blue circles) based on a clinical outcome. These compounds are run on two assays (Fig. 2A and 2B) and the measured values are obtained (e.g. XC_50_ values). Both assays are predictive as they (imperfectly) separate the toxic/safe groups, and the vertical dashed lines are optimal thresholds that minimise the misclassification error. If a compound is to the right of the dashed line it is considered “positive” in the assay and given a value of 1, otherwise a value of 0. Combining the assay results by summing these values results in a total score of either 0, 1, or 2. Compounds with a score of 2 fall in the top right quadrant of Figure 2C. These compounds are all toxic, but many toxic compounds are in other quadrants. Compounds with a score of at least 1 fall into the top left, top right, and bottom right quadrants. Although all the toxic compounds are in these three quadrants, so are some safe compounds.

**Figure 2.**
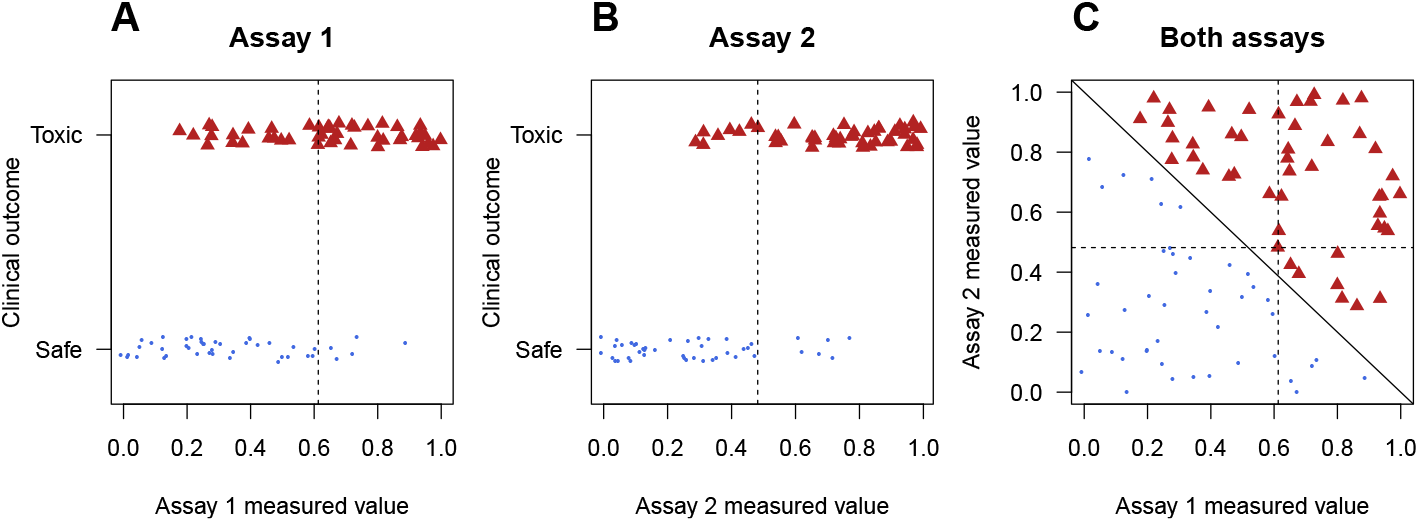
The bin-and-sum method misses simple patterns. Two assays predict a clinical outcome (A, B), but summing the number of positive assay results for each compound provides poor predictions. These compounds can be perfectly separated with a statistical model, represented by the diagonal solid line (C). Dashed lines are the optimal univariate thresholds from panel A and B.

Table 1 reports the sensitivity, specificity, and accuracy of each assay and their combination using two rules: call a compound “toxic” if it is positive in at least one assay, or call a compound “toxic” if it is positive in both assays. This approach is sub-optimal compared with fitting a statistical model to the data, which perfectly classifies the compounds, indicted by the solid diagonal line (Fig. 2C). The dashed lines are the same optimal univariate thresholds obtained from Figure 2A and 2B.

**Table 1.** Both assays predict the clinical outcome but their combination is not as effective as using a model.

| Predictor | Sensitivity | Specificity | Accuracy |
| --- | --- | --- | --- |
| Assay 1 | 0.86 | 0.70 | 0.76 |
| Assay 2 | 0.87 | 0.84 | 0.86 |
| Either | 0.81 | 1 | 0.88 |
| Both | 1 | 0.64 | 0.74 |
| Model | 1 | 1 | 1 |

The final problem with the bin-and-sum method is that it cannot capture cases when two or more weak signals are highly predictive of toxicity. Alternatively, values for two or more assays may be correlated because they are measuring a similar process or mechanism, such as cell death. Compounds will tend to be positive or negative on all the correlated assays, leading to “double-” or “triple-counting”. Both of these effects can be captured with interaction terms in a statistical model.

### Methods to prevent overfitting are inadequate

Overfitting occurs when a prediction model or rule is too flexible and accounts for idiosyncratic aspects of the data that will not be observed in future data. An overfit model will predict the current data well but will predict future data poorly, and it is very easy to overfit. Figure 3 illustrates overfitting, where data were simulated for 100 compounds, 50 toxic and 50 safe. Four predictor variables (x1 to x4) were generated from a uniform distribution and have no predictive value. Even so, we can build a decision tree with 70% accuracy using the following decision rules (Fig. 3A). First, if a compound has a value for x3 less than 0.17, classify it as toxic (right branch). 14 out of 18 compounds with x3 < 0.17 are toxic. If a compound has a value for x3 < 0.17 then variable x1 is used for the next decision. If x1 < 0.9 then a compound is classified as safe, and all 7 compounds in this far-left branch are correctly classified. And so on for the other decision points. This tree has several constraints to minimise the complexity, otherwise, we can continue creating decision points until all the compounds are correctly classified. The constraints are: (1) a maximum of four splits can be made (potentially one for each variable), a split was not made if there were fewer than 20 compounds in a branch, and a split was not made if it leads to a final node with less than five compounds. Despite these constraints, we obtain 70% accuracy when the predictor variables are random noise.

**Figure 3.**
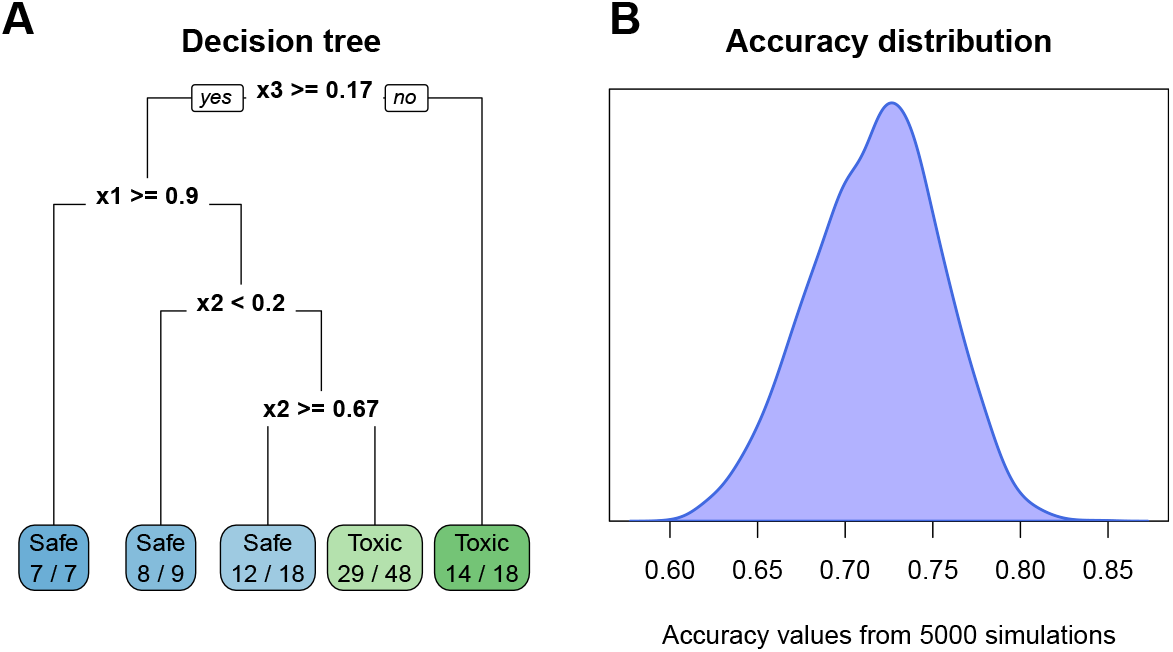
Overfitting. A decision tree predicts a toxic or safe outcome using four variables *(x* 1 to x4) that are pure noise with 70% accuracy (A). Simulating this process with 5000 datasets gives a distribution of accuracy values with a mean of 72% and a range of 60% to 85% (B).

To better appreciate how easy it is to get a respectable prediction with pure noise, 5000 datasets were generated and analysed as above (although a different tree was constructed for each dataset using the same process), and the accuracy values plotted in Figure 3B. The average accuracy was 72%, with a range of 60% to 85%.

A common way to prevent overfitting is to split the data into a training set that is used to develop a model and a test set that is used to evaluate the model. When building a model, many analytical decisions must be made and values for parameters that are not learned from the data must be selected. Cross-validation and bootstrapping the training data are methods that reduce the chance of overfitting [21]. However, it is possible for the independence of the training and test data to breakdown when multiple models are fit to the training data and the performance on the test data informally feeds back to select a model or tune the hyperparameters. This may be difficult to avoid when results are presented to a project team and members suggest to modify the model; for example, to include additional variables. After several rounds of such a process, the performance on the test set can be highly optimistic. Therefore it is often beneficial to have a third dataset that is only used to evaluate the performance of the final model, although with the small size of many safety pharmacology datasets, this is often unrealistic.

A benefit of the recent interest in machine learning and artificial intelligence is a greater awareness of assessing predictive performance on an independent test set. If standard machine learning methods are used for these prediction problems then workflows will include an assessment of overfitting. However, many prediction models in industry are still hand-crafted or created by those with little training in predictive modelling.

### Ignoring uncertainty

Uncertainty exists in many aspects of predicting, and for most toxicology and safety pharmacology applications, uncertainty needs to be incorporated into models and reported with the final predictions. For example, if a climate model predicts a 2°C increase in global temperature in the next 20 years, one’s interpretation differs if the uncertainty lies between 1.8 to 2.2°C versus −8 to 12°C. In the latter case the uncertainty is so great that no sensible actions can be taken (should we prevent global warming or global cooling?).

As mentioned above, the data are a source of uncertainty; assay values are often measured with error and some predictor variables may themselves be predictions, such as C_max_ values. Parameters in statistical models (*β*’s in Equations 2 and 3 above) are also uncertain. Since predictions from such models are based on the *β* values, greater uncertainty in the parameters will lead to greater uncertainty in the predictions. As the sample size increases, uncertainty in parameters decreases, but more samples enables more complex models to be fit, and so parameter uncertainty rarely reduces to zero in practice, even with a large sample size.

The models used to make predictions are also uncertain; we do not have a “true” model that we take as given and use for predictions. Many models may provide similar predictions, with no single model dominating. For example, if we have ten x variables (assays, physical/chemical properties, C_max_, etc.) in Equation 3, which combination of these variables should we use in a regression model? Do we include any interactions to allow for the multiplicative effect of weak signals, and if so, between which predictors? Are all the relationships linear between the x’s and the outcome, or do we expect some “U” or inverted-U relationships? We also need not restrict ourselves to a regression model but can also consider random forests, support vector machines, or neural network models. Although many predictive models can be created, predictions are typically only made from one model and thus model uncertainty is ignored. Model uncertainty can be incorporated into predictions using ensembles, stacking, or model-averaging [22, 23].

Finally, even if all the above sources of uncertainty are reduced to near zero, the predictions themselves will still be uncertain because we can rarely make perfect predictions of clinical outcomes from preclinical assays and compound properties. Unfortunately, many prediction models only report a point prediction or the “best estimate” that is then used for decision making. Bayesian methods provide a principled approach to quantify, integrate, and report uncertainty in predictions, and are now used in production for cardio and liver toxicity [5, 6], and are being developed for other applications [24, 25, 26, 27, 28, 29]. Note that we are not referring to naive Bayes classifiers, which do not place priors on all unknowns and update them with data [30], but rather full Bayesian versions of standard statistical or machine learning methods, such as logistic regression and neural networks, as well as Gaussian processes and Bayesian Additive Regression Trees (BART).

### Narrowly focusing on a few metrics

After a model is developed, its performance is assessed, but the metrics commonly examined are limited. The discussion below mainly applies to classification tasks, where the aim is to predict a binary outcome (safe/toxic), but is also relevant for categorical outcomes (safe/apoptosis/necrosis) and ordered categorical outcomes (safe/mild/severe). Four types of metrics should be examined: classification, discrimination, calibration, and overall fit.

#### Classification

metrics are the most common and include accuracy, sensitivity, and specificity. Although easy to interpret, they can be misleading when the classes are unbalanced. For example, if 90% of compounds are non-toxic, we can achieve 90% accuracy by ignoring any experimental data and always predicting “safe”. A better option is to report balanced accuracy, which is the average accuracy of the two classes. Table 2 shows the result of a hypothetical prediction model. The standard accuracy calculation is 92%:

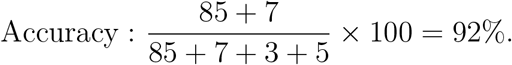

Such a high accuracy may appear impressive, but we can get 90% accuracy by always predicting a compound is safe. The balanced accuracy is only 82%, which indicates that the model is worse than always predicting the most frequent class, at least based on the accuracy criterion:

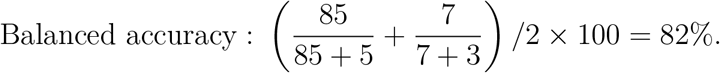

**Table 2.** Accuracy and balanced accuracy.

|  |  | Actual |  |  |
| --- | --- | --- | --- | --- |
| Predicted |  | Safe | Toxic | Total |
|  | Safe | 85 | 3 | **88** |
|  | Toxic | 5 | 7 | **12** |
| Total |  | **90** | **10** | **100** |

Assays and models are often ranked on their accuracy, sensitivity, and specificity, but with small datasets these metrics have low precision. For example, given the data in Table 2, the 95% confidence interval (CI) for standard accuracy is 84.8% to 96.5%, making assays or models that differ in accuracy by as much as 10% difficult to distinguish. Most statistical software can calculate CIs as these metrics are based on a binomial proportion. Reporting CIs can prevent over-interpreting small differences between assays or models.

A second problem is that to calculate these metrics we have converted a continuous prediction probability into a class using a threshold that may not be meaningful. A logical threshold is 0.5, with compounds classified as toxic if the prediction is > 0.5, and safe if the prediction is ≤ 0.5. However, another threshold might better reflect the costs of misclassifying a safe compound as toxic, and vice versa. Furthermore, this is another example of losing information by binning. Many predictive methods such as logistic regression, naive Bayes, and neural networks provide a continuous value for the probability of toxicity, and models are better assessed on this. Consider two models (or people) that predict a compound’s toxicity with probabilities 0.55 and 0.95. If the compound is toxic, and taking 0.5 as a threshold for toxicity, both models made correct predictions, but the second made a stronger prediction and should therefore be considered better. Similarly, if the drug is safe, the model that predicted 0.95 should be penalised more than the model that predicted 0.55. Both models made wrong predictions, but one was more wrong than the other. Methods that assess the quality of a prediction are known as scoring rules or scoring functions, and good scoring rules distinguish between the above situations [31]. Accuracy fails, but so do sensitivity and specificity, which also depend on the balance of the classes and require converting a continuous probability into a binary category. Furthermore, sensitivity and specificity do not tell us what we want to know when making a prediction for a new compound, which is the probability that a compound is safe, given that it is predicted to be safe: P(safe | predicted safe). Instead they tell us the probability that a compound is predicted to be safe, given that it is actually safe: P(predicted safe| actually safe) [32]. Although classification metrics tend to receive considerable attention, they do not tell the full story.

#### Discrimination

is the ability of a model to distinguish between classes. A common measure is the concordance index (c-index), which equals the area under the receiver-operating curve (AUC) when the variable predicted has two classes [33]. The c-index is a number between 0 and 1 giving the probability that a randomly selected toxic compound has a predicted score higher than a randomly selected nontoxic compound. Perfect discrimination gives a c-index of 1 and a model with no predictive ability will give a value of 0.5. The c-index requires no arbitrary threshold and is unaffected by class imbalances. We use data from Pollard et al. to illustrate discrimination and the other metrics [34]. This dataset is used to predict QT prolongation, a binary (Yes = 1 / No = 0) outcome based on clinical C_max_ values and IC_50_ values from a hERG inhibition assay. Further details on the data and model can be found in Lazic et al. [5], and in the supplementary material. The c-index for the model equals 0.94, and for comparison, accuracy = 89.7%, sensitivity = 90.9%, and specificity = 88.9%. Although uncommon, the c-index can be represented as a number between 0-100% if the other metrics are also on a percentage scale.

#### Calibration

measures the agreement between the predictions and true outcomes. A model is calibrated when the predicted probabilities match the observed probabilities, for all values of the predicted probabilities. Even if a model has reasonable accuracy, it can still over- or under-predict the true probabilities, and this is often assessed graphically. Figure 4A plots the predicted probabilities on the *x*-axis and the observed proportions on the *y*-axis, with calibrated predictions falling along the diagonal red line. The curved blue line shows the results from the model and the grey band indicates the 95% confidence interval. The blue line is close to the red, but with few compounds, substantial uncertainty is present. Calibration curves are discussed in [32, 21].

**Figure 4.**
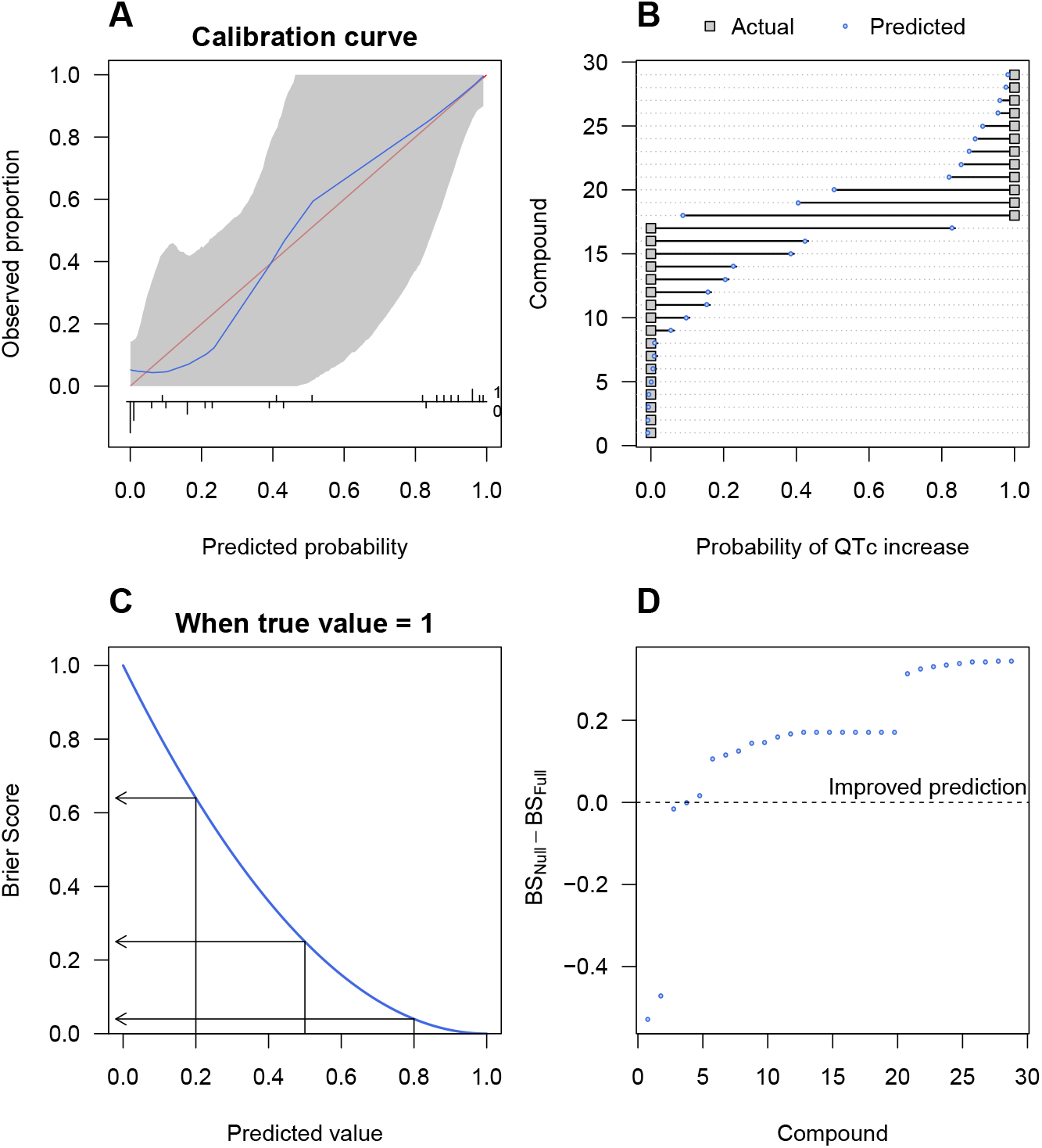
Model evaluation metrics. A calibration curve plots the predicted versus actual outcomes, and a well-calibrated model (blue line) will be close to the red diagonal line (A). The Brier score (BS) is calculated as the squared difference between the actual and predicted values (B). The relationship between predictions and BS when the true class is 1 (C). The difference in BS between a null and full model shows the improvement in BS for most compounds (D).

Machine learning algorithms are trained by maximising the *overall fit* of a model to the data. Some metrics of overall fit are difficult to interpret and therefore infrequently reported. The Brier score (BS) has a simple interpretation and is appropriate when models return a probability for a binary outcome [35] (see Semenova et al. for an extension to ordered categorical outcomes [29]). The BS is the squared difference between the prediction – a value between 0 and 1 – and the actual category, which is either 0 or 1:

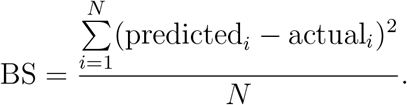

*N* is the number of compounds and *i* indexes the compounds. The greater the difference between the actual and predicted values, the larger the BS, and so low scores are better. We get a BS for each compound, and by summing all the scores and dividing by *N* we get an average BS for the dataset. The grey squares in Figure 4B are the true values for QT prolongation from the Pollard et al. data and the blue circles are the predicted values [34]. The square of the length of the line connecting the two points is the BS for that compound.

Figure 4D shows the difference in BS for two models. The first is a “null” model that does not contain any predictor variables. The model can still achieve a 59% prediction accuracy because 59% of compounds do not increase the QT interval. The second model uses C_max_ and hERG IC_50_ and is called the “full” model. Since the BS is expected to be higher for the null model (worse predictions), positive values for the difference BS_Null_ – BS_Full_ indicate compounds that are better predicted when using C_max_ and hERG IC_50_, compared with not using this information. The average BS for the null model is 0.24 and for the full model is 0.09, indicating a large improvement. Even though the prediction for three compounds is worse when including the C_max_ and hERG data, the average performance is better. The average BS estimates the overall model fit, and the individual Brier scores indicate which compounds are hard to predict. These can be further investigated to understand why the model was unable to predict their outcome. For example, they may inhibit other cardiac ion channels.

The BS can be scaled between 0 and 1 using [36]:

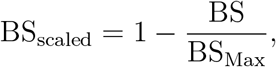

where BS_Max_ is the largest BS possible when always predicting the most frequent class, and averaged over all of the compounds (the null model). A scaled BS of 0 means we predict no better than always choosing the most frequent class, and a value of 1 means we always predict the true class with maximum probability. It can also be interpreted as an *R^2^* value, or the proportion of variance in the outcome that is explained by the predictors [37]. For this example, BSscaled = 0.62.

The commonly reported metrics of accuracy, sensitivity, and specificity have limitations and do not fully describe the performance of a prediction model. Hence, we suggest also examining discrimination, calibration, and overall model fit metrics when making decisions [38].

### Not assessing the domain of applicability

Once a suitable prediction model has been built and is put into production, it can still perform poorly if the new compounds differ from those used to build the model. A final problem therefore is that the similarity between new (test) compounds and old (training) compounds is often not assessed; in other words, the model may be applied outside of its relevant domain. New compounds may differ structurally from the training compounds as new chemical space is explored, resulting in different physiochemical properties and assay results. If new compounds lie outside of the training data in multidimensional space, prediction becomes extrapolation – a dangerous procedure. Alternatively, new compounds may be located within the training data, but in regions with few observations. Even though the model is now interpolating, predictions may be poor when data are sparse. Some models such as Gaussian Processes can account for both of these situations by allowing the uncertainty in the prediction to increase, but this is not possible for most standard machine learning models [39].

Assessing similarity between old and new compounds should therefore be routine when making predictions. Many methods are available to detect if a data point is unlike others, known as outlier detection or anomaly detection [40]. To illustrate this point and one solution, the hERG IC_50_ and C_max_ data and the model from the previous section are used [5]. The idea is to characterise the hERG IC_50_ and C_max_ values with a mixture of Gaussian distributions, which is a form of clustering or unsupervised machine learning [41]. Then for a new compound, we calculate its distance to the nearest cluster. If this distance is large, we can flag the compound as an outlier and be more suspicious of the prediction. We use the Mahalanobis distance, which takes the shape and orientation of the clusters into account when calculating the distance to the cluster centre.

Figure 5A plots the hERG IC_50_ and C_max_ training data (black circles) and the shaded blue regions show the three clusters that describe the data. The optimal number of clusters was determined by fitting 1-4 Gaussian distributions to the data and assessing their fit while penalising overly complex descriptions. We do not interpret the clusters, and only use them to describe the structure of the data. The four red diamonds labelled A–D are the hypothetical new compounds and were not used to form the clusters.

**Figure 5.**
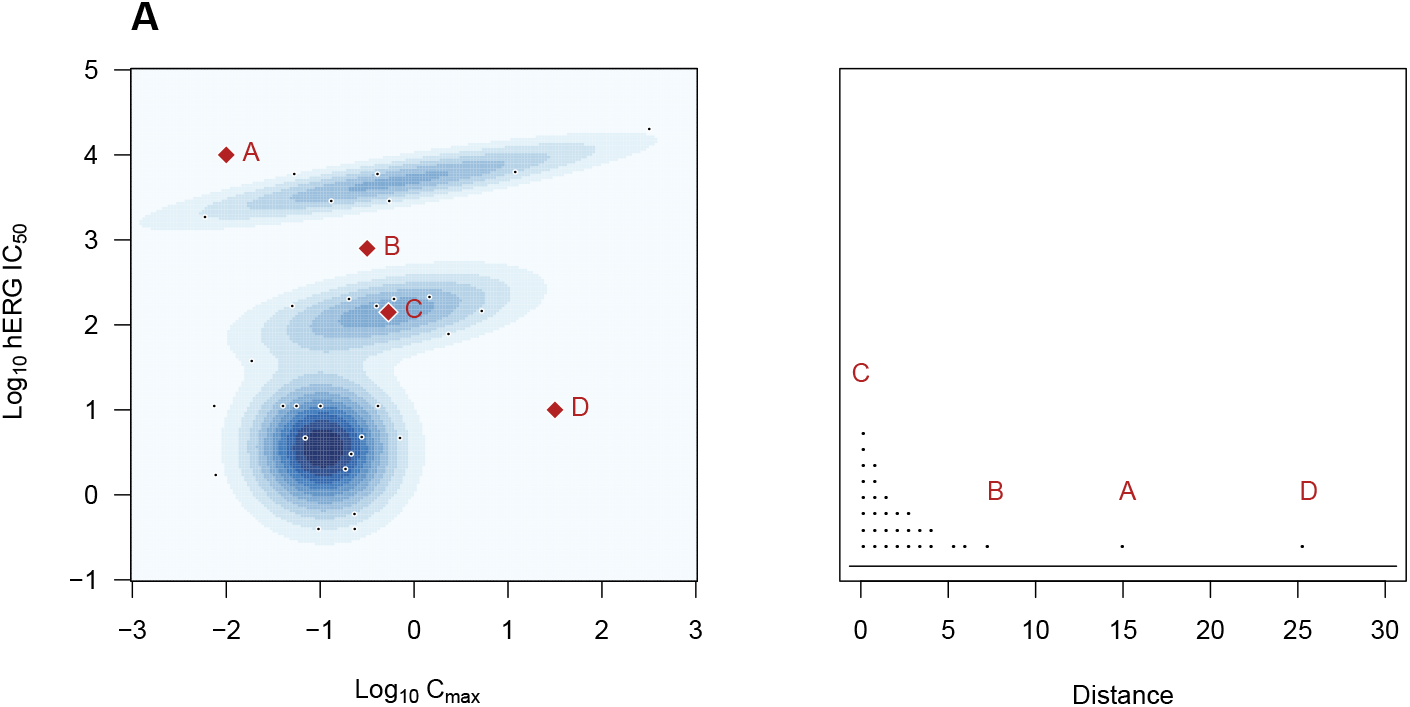
Outlier detection. The training data (black circles) are described by a mixture of three Gaussian distributions (shaded regions; A). The distance of four new data points (red diamonds, A–D) to the nearest cluster centre are calculated (B). Compounds with a large distance are unlikely to be from one of the clusters and their predictions may be unreliable.

Figure 5B plots the distance to the nearest cluster centre for the original and the four new compounds. Compound D is furthest from the original data in Figure 5A and has the largest distance. Note that compound D does not have the lowest hERG IC_50_ value nor the largest C_max_ value, and so looking at one variable at a time would not indicate that compound D is unusual. Compound A is closer to the bulk of the data but still near the edge, and thus has a large distance. Compound B lies within the interior of the data but in a region with few observations. It therefore has a larger distance. Compound C lies at the mean of the middle cluster and therefore has the smallest distance of all the compounds, both old and new (although this cannot be seen in Fig. 5B due to the small amount of binning required to stack the data points).

With only two predictor variables, visualising an unusual data point is simple, but it is much more difficult in higher dimensions. Formal methods to flag unusual compounds are therefore useful to highlight compounds for which predictions may be unreliable [42, 43, 40].

## Discussion and recommendations

Why do poor practices persist? First, inertia: methods become established as standard operating procedures in a field or company, despite their shortcomings, and they are not updated when additional limitations are discovered years later. Second, many of the above practices are simple to use and easy to understand. Many people who develop and use these predictive models may not have a statistics, data science, or machine learning background, and therefore rely on simpler but less suitable methods. Finally, there is little learning from failure at the institutional level in pharmaceutical companies, where these models are developed and used. A long lag exists between making a preclinical decision and learning the eventual clinical outcome. Often years pass, and with staff turnover, few of the original decision makers may be around to reflect on the choices or methods used. By documenting the decision and reasons for it – ideally in a machine-readable format – this information can be fed back to improve decisions years later.

We discussed several common practices that can lead to problems when predicting toxicity. These problems happen at all stages of model development, including preprocessing (converting continuous variables into binary variables using arbitrary thresholds), feature engineering (using margins), model building (using scores, overfitting, ignoring uncertainty), assessment (evaluation metrics), and production (similarity of old and new compounds). This paper provides a high-level summary of current practices and the references cited throughout provide more theoretical background and technical details on implementing these methods. In addition, the data and R-code are provided on Github to facilitate the uptake of these methods (https://github.com/stanlazic/toxdata_2020).

Many recommendations are simple to apply: avoid binning variables, use the measured continuous values instead; avoid using margins in predictive models, use IC_50_ and C_max_ values directly; and when evaluating the suitability of a predictive model, consider four types of metrics: classification, discrimination, calibration, and overall fit. If classes are unbalanced, prefer balanced accuracy to overall accuracy. Other recommendations will require a collaboration with quantitative researchers, such as avoiding hand-crafted arbitrary scores and using statistical or machine learning models to predict outcomes. Similarly, assessing the domain of applicability is more difficult as many options are available, but any method is better than none. Quantifying and propagating uncertainty is a large topic that was only briefly mentioned, but Bayesian methods are the standard way of handling uncertainty and are becoming more popular for predictive models in safety pharmacology [24, 25, 26, 27, 28, 29]. Large and small pharmaceutical and biotechnology companies, contract research organisations, and other industries involved in drug development need to understand the limitations of using these assays out of context. A robust statistical framework is required to maximise predictivity, prevent errors of judgement, and reduce cost from incorrect use.

