## Supplementary material for "Improving drug safety predictions by reducing poor analytical practices": Data and code: code.html (from another version of this preprint: version 1, 2020-09-28)

###### Stanley E. Lazic

Load packages and functions.

```
knitr::opts_chunk$set(message = FALSE)
```

```
library(tidyverse)
library(pROC)
library(rpart)
library(rpart.plot)
library(patchwork)
library(CalibrationCurves)
library(mclust)
library(caret)

source("functions.R") # collection of user-defined functions
```

### Use of margins is misleading

Generate data for two assays (x1, x2) and C_max_ for 100 compounds.

```
set.seed(123)
x1 <- runif(100, 1, 2)
x2 <- runif(100, 1, 2)
cmax <- runif(100, 1, 2)
```

Generate values for the outcome (y), which depends only on C_max_, and calculate the safety margins for the assays.

```
y <- cmax + rnorm(100, 0, 0.15)
x1_marg <- x1 / cmax
x2_marg <- x2 / cmax
```

Combine all the values into a dataframe and plot. Note how the assays are uncorrelated with the outcome and therefore have no predictive value. The margins however do correlate with the outcome and they appear to be good predictors, but these relationships are completely driven by C_max_ -- the assays are still useless. Furthermore, the margins are now correlated with each other, which can be a problem for some predictive models.

```
d <- data.frame(y, x1, x2, cmax, x1_marg, x2_marg)

p1 <- ggplot(aes(x1, y), data = d) +
    geom_point(col = "royalblue", size = 1.75) + ylab("Outcome") +
    xlab("Assay 1") + theme_classic(base_size = 16)

p2 <- ggplot(aes(x2, y), data = d) +
    geom_point(col = "royalblue", size = 1.75) + ylab("Outcome") +
    xlab("Assay 2") + theme_classic(base_size = 16)

p3 <- ggplot(aes(cmax, y), data = d) +
    geom_point(col = "royalblue", size = 1.75) + ylab("Outcome") +
    xlab("Cmax") + theme_classic(base_size = 16)

p4 <- ggplot(aes(x1_marg, y), data = d) +
    geom_point(col = "royalblue", size = 1.75) + ylab("Outcome") +
    xlab("Margin (Assay 1/Cmax)") + theme_classic(base_size = 16)

p5 <- ggplot(aes(x2_marg, y), data = d) +
    geom_point(col = "royalblue", size = 1.75) + ylab("Outcome") +
    xlab("Margin (Assay 2/Cmax)") + theme_classic(base_size = 16)

p6 <- ggplot(aes(x1_marg, x2_marg), data = d) +
    geom_point(col = "royalblue", size = 1.75) + ylab("Margin (Assay 2/Cmax)") +
    xlab("Margin (Assay 1/Cmax)") + theme_classic(base_size = 16)

p1 + p2 + p3 + p4 + p5 + p6 + plot_annotation(tag_levels = "A")
```

### Problems with binning and thresholds

Generate data for 100 compounds for two assays (x1, x2) and a variable indicating if the compounds are toxic (group). If the sum of the assay values is greater than 1, the compound is toxic (group=1), otherwise it is safe (group=0).

```
set.seed(8)
x1 <- runif(100)
x2 <- runif(100)
group <- ifelse(x1 + x2 < 1, 0, 1)
```

Remove compounds whose sum of assay values is between 0.95 and 1.05. This creates a gap between the groups in 2-dimensional space to separate the groups (for visual effect).

```
keep_index <- ifelse(x1 + x2 > 0.95 & x1 + x2 < 1.05, 0, 1)

x1 <- x1[keep_index == 1]
x2 <- x2[keep_index == 1]
group <- group[keep_index == 1]
N <- length(group) # number of remaining compounds
```

Calculate the optimal thresholds that separates the groups for each assay.

```
## calculate optimal thresholds
roc1 <- roc(group ~ x1)
roc2 <- roc(group ~ x2)

## extract threshold values for plotting
coords(roc1, x = "best", input = "threshold", transpose = FALSE) %>%
    .[, "threshold"] ->
    threshold1

coords(roc2, x = "best", input = "threshold", transpose = FALSE) %>%
    .[, "threshold"] ->
    threshold2
```

Plot the resulting data and thresholds.

```
## specify colours and plotting symbols
pchs <- ifelse(group == 0, 16, 17)
cols <- ifelse(group == 0, "royalblue", "firebrick")

par(mfrow = c(1, 3),
    mar = c(4.5, 3.8, 3, 1),
    las = 1,
    cex = 1.2)

plot(jitter(group, factor = 0.3) ~ x1, yaxt = "n",
     ylab = "Clinical outcome", xlab = "Assay 1 measured value", main = "Assay 1",
     ylim = c(-0.2, 1.2), pch = pchs, col = cols)
axis(2, at = 0:1, labels = c("Safe", "Toxic"))
abline(v = threshold1, lty = 2)
mtext("A", side = 3, line = 1.5, font = 2, cex = 2, adj = 0)

plot(jitter(group, factor = 0.3) ~ x2, yaxt = "n",
     ylab = "Clinical outcome", xlab = "Assay 2 measured value", main = "Assay 2",
     ylim = c(-0.2, 1.2), pch = pchs, col = cols)
axis(2, at = 0:1, labels = c("Safe", "Toxic"))
abline(v = threshold2, lty = 2)
mtext("B", side = 3, line = 1.5, font = 2, cex = 2, adj = 0)

plot(x2 ~ x1,
     pch = pchs, col = cols, ylab = "Assay 2 measured value",
     xlab = "Assay 1 measured value", main = "Both assays",
     ylim = c(0, 1), xlim = c(0, 1))
abline(v = threshold1, h = threshold2, lty = 2)
abline(1, -1)
mtext("C", side = 3, line = 1.5, font = 2, cex = 2, adj = 0)
```

Calculate metrics for the first assay.

```
##
confusionMatrix(data = factor(group),
                reference = as.factor(as.numeric(x1 > threshold1)),
                positive = "1")
```

```
## Confusion Matrix and Statistics
## 
##           Reference
## Prediction  0  1
##          0 39  5
##          1 17 31
##                                         
##                Accuracy : 0.761         
##                  95% CI : (0.661, 0.844)
##     No Information Rate : 0.609         
##     P-Value [Acc > NIR] : 0.0015        
##                                         
##                   Kappa : 0.526         
##                                         
##  Mcnemar's Test P-Value : 0.0190        
##                                         
##             Sensitivity : 0.861         
##             Specificity : 0.696         
##          Pos Pred Value : 0.646         
##          Neg Pred Value : 0.886         
##              Prevalence : 0.391         
##          Detection Rate : 0.337         
##    Detection Prevalence : 0.522         
##       Balanced Accuracy : 0.779         
##                                         
##        'Positive' Class : 1             
##
```

Calculate metrics for the second assay.

```
confusionMatrix(data = factor(group),
                reference = as.factor(as.numeric(x2 > threshold2)),
                positive = "1")
```

```
## Confusion Matrix and Statistics
## 
##           Reference
## Prediction  0  1
##          0 38  6
##          1  7 41
##                                        
##                Accuracy : 0.859        
##                  95% CI : (0.77, 0.923)
##     No Information Rate : 0.511        
##     P-Value [Acc > NIR] : 2.22e-12     
##                                        
##                   Kappa : 0.717        
##                                        
##  Mcnemar's Test P-Value : 1            
##                                        
##             Sensitivity : 0.872        
##             Specificity : 0.844        
##          Pos Pred Value : 0.854        
##          Neg Pred Value : 0.864        
##              Prevalence : 0.511        
##          Detection Rate : 0.446        
##    Detection Prevalence : 0.522        
##       Balanced Accuracy : 0.858        
##                                        
##        'Positive' Class : 1            
##
```

Calculate a "score" for each compound by summing the number of assays the compound was positive in. Calculate the metrics for two decision rules: (1) call a compound toxic if it is positive in at least one assay, or (2) call a compound toxic if it is positive in both assays.

```
## caclulate "score"
score <- as.numeric(x1 > threshold1) + as.numeric(x2 > threshold2)

## positive in either assay
confusionMatrix(data = factor(group),
                reference = as.factor(as.numeric(score >= 1)),
                positive = "1")
```

```
## Confusion Matrix and Statistics
## 
##           Reference
## Prediction  0  1
##          0 33 11
##          1  0 48
##                                         
##                Accuracy : 0.88          
##                  95% CI : (0.796, 0.939)
##     No Information Rate : 0.641         
##     P-Value [Acc > NIR] : 2.09e-07      
##                                         
##                   Kappa : 0.758         
##                                         
##  Mcnemar's Test P-Value : 0.00257       
##                                         
##             Sensitivity : 0.814         
##             Specificity : 1.000         
##          Pos Pred Value : 1.000         
##          Neg Pred Value : 0.750         
##              Prevalence : 0.641         
##          Detection Rate : 0.522         
##    Detection Prevalence : 0.522         
##       Balanced Accuracy : 0.907         
##                                         
##        'Positive' Class : 1             
##
```

```
## positive in both assays
confusionMatrix(data = factor(group),
                reference = as.factor(as.numeric(score == 2)),
                positive = "1")
```

```
## Confusion Matrix and Statistics
## 
##           Reference
## Prediction  0  1
##          0 44  0
##          1 24 24
##                                         
##                Accuracy : 0.739         
##                  95% CI : (0.637, 0.825)
##     No Information Rate : 0.739         
##     P-Value [Acc > NIR] : 0.555         
##                                         
##                   Kappa : 0.489         
##                                         
##  Mcnemar's Test P-Value : 2.67e-06      
##                                         
##             Sensitivity : 1.000         
##             Specificity : 0.647         
##          Pos Pred Value : 0.500         
##          Neg Pred Value : 1.000         
##              Prevalence : 0.261         
##          Detection Rate : 0.261         
##    Detection Prevalence : 0.522         
##       Balanced Accuracy : 0.824         
##                                         
##        'Positive' Class : 1             
##
```

Combining the assays by summing the number of times compounds are positive in an assay to generate a score does not enable us to perfectly classify the compounds. But the compounds are perfectly separable (Panel C above). The conclusion is that the "bin-and-sum" algorithm is unsuitable.

### Overfitting

Generate data for 100 compounds for four assays (x1-x4), and an outcome (y) that is independent of the assays; that is, the assays have no predictive value.

```
set.seed(123)
x1 <- runif(100)
x2 <- runif(100)
x3 <- runif(100)
x4 <- runif(100)
y <- as.factor(rep(c("Safe", "Toxic"), each = 50))

d <- data.frame(y, x1, x2, x3, x4)
```

Use a decision tree to derive a set of rules for classifying the compounds. Restrict the tree to have no more than four splits (`maxdepth=4`), with a minimum of 20 compounds remaining before a spilt is attempted (`minsplit=20`), and a minimum of five compounds in each terminal node (`minbucket=5`). These restrictions prevent the tree from becoming too large and complicated, which would increase the overfitting even more.

```
## fit decision tree
mod <- rpart(y ~ ., data = d, method = "class",
             control = list(maxdepth = 4, minsplit = 20, minbucket = 5, xval = 0))

## calculate the overall accuracy
xtabs(~ y + predict(mod, type = "class")) %>%
    diag() %>%
    sum() / 100 ->
    acc
```

The accuracy is 70%, which is high considering the assays were generated to have zero predictive ability. Next, we repeat the above process 5000 times and examine the accuracy values (the function `overfit()` is defined in the `functions.R` file and wraps the above code to easily repeat the data generation and calculation.)

```
## simulate 5000 datasets
set.seed(123)
acc_sims <- replicate(5000, overfit())

## caclulate summary statistics
summary(acc_sims)
```

```
##    Min. 1st Qu.  Median    Mean 3rd Qu.    Max. 
##   0.600   0.690   0.720   0.717   0.740   0.850
```

The mean accuracy for the 5000 simulations is 71.7% with a range from 60% to 85%. Below we plot the first decision tree and a density plot of the accuracy values from the simulations.

```
## calcualte values for density plot
dens <- density(acc_sims, adjust = 1.25)

par(mfrow = c(1, 2),
    mar = c(4.5, 1, 3, 0.5),
    las = 1,
    cex = 1.2)

rpart.plot(mod, type = 0, extra = 2)
title("Decision tree")
mtext("A", side = 3, line = 1.5, font = 2, cex = 2, adj = 0)

plot(dens, main = "Accuracy distribution", ylab = "", yaxt = "n", yaxs = "i",
     xlab = "Accuracy values from 5000 simulations", ylim = c(0, 11))
polygon(dens, border = "royalblue", col = rgb(0, 0, 1, 0.3), lwd = 2)
mtext("B", side = 3, line = 1.5, font = 2, cex = 2, adj = 0)
```

### Brier score, calibration, and discrimination

Load cardiotoxicity data and remove two compounds that work through a different mechanism.

```
d <- read.delim("QT_data.txt")

# remove beta2 agonists
d <- d[!d$Compound %in% c(9, 12), ]
```

Fit two models to the data: the first (`m1`) has no predictor variables, the second (`m2`) predicts whether a compound increases the QT interval (an indicator of cardiotoxicity) using Log10 C_max_ and the Log10 hERG IC50 value from an inhibition assay. The model predictions are extracted and the Brier scores calculated.

```
m_null <- glm(QT.increase ~ 1, data = d, family = "binomial")
m_full <- glm(QT.increase ~ log.cmax * log.herg, data = d, family = "binomial")

pred_null <- invlogit(predict(m_null))
pred_full <- invlogit(predict(m_full))

bs_null <- brier_score(pred_null, d$QT.increase)
bs_full <- brier_score(pred_full, d$QT.increase)

## calculate the mean Brier score
mean(bs_null)
```

```
## [1] 0.242568
```

```
mean(bs_full)
```

```
## [1] 0.0927506
```

```
## calculate the scaled Brier score for the model
brier_score(pred_full, d$QT.increase, scale = TRUE) %>% mean()
```

```
## [1] 0.617631
```

Calculate the standard accuracy, sensitivity, and specificity metrics.

```
confusionMatrix(data = factor(d$QT.increase),
                reference = as.factor(as.numeric(pred_full > 0.5)),
                positive = "1")
```

```
## Confusion Matrix and Statistics
## 
##           Reference
## Prediction  0  1
##          0 16  1
##          1  2 10
##                                         
##                Accuracy : 0.897         
##                  95% CI : (0.726, 0.978)
##     No Information Rate : 0.621         
##     P-Value [Acc > NIR] : 0.000989      
##                                         
##                   Kappa : 0.784         
##                                         
##  Mcnemar's Test P-Value : 1.000000      
##                                         
##             Sensitivity : 0.909         
##             Specificity : 0.889         
##          Pos Pred Value : 0.833         
##          Neg Pred Value : 0.941         
##              Prevalence : 0.379         
##          Detection Rate : 0.345         
##    Detection Prevalence : 0.414         
##       Balanced Accuracy : 0.899         
##                                         
##        'Positive' Class : 1             
##
```

Plot the calibration curve and some plots to understand the Brier scores (details in the manuscript).

```
par(mfrow = c(2, 2),
    mar = c(4.5, 4.3, 2.6, 0.5),
    las = 1,
    cex = 1.2)

val.prob.ci.2(pred_full, y = d$QT.increase, logit = "p", CL.BT = TRUE,
              dostats = FALSE, legendloc = FALSE,
              col.smooth = "royalblue", lty.smooth = 1.5)
mtext("A", side = 3, line = 1, font = 2, cex = 1.75, adj = 0)
title("Calibration curve", line = 0.5)


## index to order the compounds by QT value and then predicted value
ind <- order(d$QT.increase, pred_full)

plot(x = d$QT.increase, y = 1:29, type = "n", xlim = c(0, 1),
     ylab = "Compound", xlab = "Probability of QTc increase")
abline(h = 1:29, lty = 3, col = "grey70")
segments(d$QT.increase[ind], 1:29, pred_full[ind], 1:29, lwd = 1.5)
points(x = d$QT.increase[ind], y = 1:29, pch = 22, col = "black", bg = "grey80")
points(x = pred_full[ind], y = 1:29, pch = 21, col = "royalblue", bg = "lightblue")

par(xpd = TRUE)
legend(x = 0.5, y = 35, legend = c("Actual", "Predicted"), pch = c(22, 21),
       col = c("black", "royalblue"), pt.bg = c("grey80", "lightblue"),
       bty = "n", xjust = 0.5, ncol = 2)
par(xpd = FALSE)
mtext("B", side = 3, line = 1, font = 2, cex = 1.75, adj = 0)

## plot Brier score for various predictions when true value
## of the outcome = 1
x <- seq(0, 1, 0.01)
bx <- brier_score(x, 1)
ind <- which(x %in% c(0.2, 0.5, 0.8))

plot(bx ~ (1 - x), type = "l", yaxs = "i", lwd = 2, col = "royalblue",
     ylab = "Brier Score", xlab = "Predicted value", main = "",
     ylim = c(0, 1.05))
title("When true value = 1", line = 0.5)

segments(x[ind], c(0, 0, 0), x[ind], bx[ind])
arrows(x[ind], bx[ind], c(-0.02, -0.02, -0.02), bx[ind],
       length = 0.1, code = 2)
mtext("C", side = 3, line = 1, font = 2, cex = 1.75, adj = 0)

## caclulate difference in Brier scores
diff <- bs_null - bs_full

plot(sort(diff), pch = 21, col = "royalblue", bg = "lightblue",
     xlab = "Compound", ylab = ~ BS[Null] - BS[Full])
abline(h = 0, lty = 2)
text(x = 30, y = 0.02, labels = "Improved prediction",
     adj = c(1, 0))
mtext("D", side = 3, line = 1, font = 2, cex = 1.75, adj = 0)
```

Calculate metrics including c-index (`C (ROC)`) for discrimination and Brier score (`Brier`), which is the same as we calculate above for `m2`: 0.09.

```
val.prob.ci.2(pred_full, y = d$QT.increase, logit = "p", CL.BT = TRUE)
```

```
## Bootstrap samples are being generated.
## 
## 
## 
## 
##  A 95% confidence interval is given for the calibration intercept, calibration slope and c-statistic.
```

```
##          Dxy      C (ROC)           R2            D     D:Chi-sq          D:p            U     U:Chi-sq          U:p 
##  8.82353e-01  9.41176e-01  7.18556e-01  7.27948e-01  2.21105e+01  2.57400e-06 -6.89655e-02 -7.10543e-15  1.00000e+00 
##            Q        Brier    Intercept        Slope         Emax Brier scaled         Eavg          ECI 
##  7.96913e-01  9.27506e-02 -1.38213e-10  1.00000e+00  4.57571e-09  6.17631e-01  3.76072e-02  2.50016e-01
```

### Outlier detection

Fit Gaussian mixture models to the cardiotoxicity data with 1-4 Gaussian distributions to find the best model of the data.

```
cl <- mclustBIC(d[, 7:8], G = 1:4)
```

```
## plot and summarise results
plot(cl)
```

```
summary(cl)
```

```
## Best BIC values:
##             EVE,3      VII,2      EVV,2
## BIC      -187.826 -190.68687 -190.99055
## BIC diff    0.000   -2.86063   -3.16431
```

The best model according to the Bayesian information criterion (BIC) has three clusters with equal volumes, different shapes, and equal orientations (the `EVE` model). Fit this model to the data.

```
mod1 <- Mclust(d[, 7:8], x = cl, G = 3, modelNames = "EVE")
```

Extract the means and covariances for the three Gaussian distributions and generate four new test data points (new compounds; red diamonds in the plot below). Then, calculate the Mahalanobis distance for each old and new data point to the mean of closest cluster and plot the results. Large values indicate that a point is an outlier.

```
## extract means and covariances
xp <- extract_parms(mod1)

## test data
test_data <- data.frame(log.herg = c(4, 2.9, 2.149, 1),
                 log.cmax = c(-2, -0.5, -0.273, 1.5))

## calculate Mahalanobis distance for training and test data
md <- calc_mahalanobis(xp, rbind(d[, 7:8], test_data))

## fit the model again for plotting
mod2 <- Mclust(d[, 8:7], x = cl, G = 3, modelNames = "EVE")


par(mfrow = c(1, 2),
    mar = c(4.5, 4.3, 2.6, 0.1),
    las = 1,
    cex = 1.2)

surfacePlot(data = d[, 8:7], parameters = mod2$parameters, ylim = c(-1, 5),
            xlim = c(-3, 3), type = "image",
            xlab = ~ Log[10] ~ C[max], ylab = ~ Log[10] ~ hERG ~ IC[50])
points(log.herg ~ log.cmax, data = d, pch = 21, col = "white", bg = "black")

points(log.herg ~ log.cmax, data = test_data, cex = 1.25, col = "white", bg = "firebrick", pch = 23)
text(LETTERS[1:4], y = test_data$log.herg, x = test_data$log.cmax,
     pos = 4, col = "firebrick")
box()
mtext("A", side = 3, line = 1, font = 2, cex = 1.75, adj = 0)


BHH2::dotPlot(apply(md, 1, min), pch = 16, xlab = "Distance",
              xlim = c(0, 30), cex = 0.8)
text(LETTERS[1:4], y = c(0.15, 0.15, 0.4, 0.15), x = apply(md[30:33, ], 1, min),
     col = "firebrick")
box()
mtext("B", side = 3, line = 1, font = 2, cex = 1.75, adj = 0)
```
